# Upstream charged and hydrophobic residues impact the timing of membrane insertion of transmembrane helices

**DOI:** 10.1101/2021.12.23.474019

**Authors:** Felix Nicolaus, Fatima Ibrahimi, Anne den Besten, Gunnar von Heijne

## Abstract

During SecYEG-mediated cotranslational insertion of membrane proteins, transmembrane helices (TMHs) first make contact with the membrane when their N-terminal end is ~45 residues away from the peptidyl transferase center. However, we recently uncovered instances where the first contact is delayed by up to ~10 residues. Here, we recapitulate these effects using a model TMH fused to two short segments from the BtuC protein: a positively charged loop and a re-entrant loop. We show that the critical residues are two Arg residues in the positively charged loop and four hydrophobic residues in the re-entrant loop. Thus, both electrostatic and hydrophobic interactions involving sequence elements that are not part of a TMH can impact the way the latter behaves during membrane insertion.

## Introduction

Nearly all helix-bundle type membrane proteins in both prokaryotic and eukaryotic cells are inserted cotranslationally into their target membrane [1,2]. While the energetics of membrane insertion of transmembrane helices (TMHs) has been extensively studied both *in vitro* and *in vivo* [3–6], it is only recently that methodologies have been developed that make it possible to follow step-by-step the membrane insertion of individual TMHs as they come out of the ribosome exit tunnel. Two methods in particular can provide this kind of information: *in vitro* real-time FRET analysis of cotranslational membrane insertion [7] and *in vivo* Force Profile Analysis (FPA) where a translational arrest peptide (AP) engineered into the polypeptide chain is used to detect the force generated on the nascent chain during membrane insertion [8,9].

In a recent study, we applied FPA to three *E. coli* inner membrane proteins: EmrE, GlpG, and BtuC [9]. For all three proteins (a total of 20 TMHs), we found that, on average, a TMH starts to generate force on the nascent chain when its N-terminal end is ~45 residues away from the polypeptide transferase center (PTC) in the large ribosome subunit. A real-time FRET study of the N-terminal TMH in another *E. coli* inner membrane protein (LepB) has shown that its N-terminal end reaches the vicinity of the entry to the SecYEG translocon channel when 40-50 residues away from the PTC [7], confirming that the increase in force is caused by the encounter of the TMH with the hydrophobic translocon/lipid environment. However, FPA further revealed that not all TMHs follow the ‘45-residue rule’, and that the fine details of the force profiles (FPs) reflect context-dependent effects that promise to provide further insights into the molecular interactions that drive and control membrane insertion of TMHs [9].

Here, we focus on two such unexpected features in the FP recorded for the 10-TMH BtuC protein [9]: (i) TMH2 starts to generate force only when its N-terminal end is ~55 rather than the expected ~45 residues away from the PTC, and (ii) two narrow peaks appear in the force profile flanking the location where a single peak generated by TMH6 was expected. In particular, we ask whether these effects are peculiar to BtuC, or whether they are also seen when the segments that flank BtuC TMH2 and TMH6 are fused to a model TMH composed of Leu and Ala residues. Indeed, we find that the effects are seen also with model TMHs, and further pin-point the implicated sequence determinants to two Arg residues located in the periplasmic loop upstream of BtuC TMH2 and a four-residue hydrophobic stretch located in a re-entrant loop upstream of BtuC TMH6. We hypothesize that the positively charged Arg residues are temporarily prevented from translocating through the ‘hydrophobic gasket’ [10] in the middle of the SecYEG channel by the membrane potential and thereby delay membrane insertion of the following TMH, and that the partitioning of the hydrophobic stretch into the periplasmic leaflet of the inner membrane somehow weakens the initial interaction between the following TMH and the lipid bilayer.

## Materials and methods

### Enzymes and chemicals

Enzymes and other reagents were purchased from Thermo Fisher Scientific, New England Biolabs, and Sigma-Aldrich. Oligonucleotides were ordered from Eurofins Genomics. L- [^35^S]-methionine was provided by PerkinElmer.

### Cloning and Mutagenesis

In order to generate the constructs used in this study the previously described pING1 plasmid harboring a shortened LepB sequence from *E. coli* with an inserted hydrophobic H-segment (4L/15A or 6L/13A) followed by a variable LepB-derived linker (between 4 and 47 residues long), the 17-residue long SecM(*Ec*) AP from *E. coli*, and a C-terminal extension containing 23 residues derived from *E. coli* LepB was employed [8]. All sequences are summarized in Supplementary Fig. S1. The DNA sequences encoding the flanking regions (RGELFVWQIRLPR; LMYWMMGGFGGVDWRQS) derived from *E. coli* BtuC (periplasmic loop connecting TMH1 and TMH2; re-entrant loop connecting TMH5 and TMH6, respectively) were amplified by PCR from a full-length LepB-BtuC-SecM(*Ec*) construct used in a previous study [9] and cloned upstream of 4L/15A or 6L/13A using Gibson assembly^®^ [11]. Site-specific DNA mutagenesis was performed to change Arg residues to Gln residues (mutants QRR, RRQ, QRQ, QQQ) in RGELFVWQIRLPR, and to replace *n* residues in LMYWMMGGFGGVDWRQS with Gly-Ser (GS) repeats (mutants *n* = 5, *n* = 9, *n* = 13, *n* = 17, and LMYW→GSGS). All cloning and mutagenesis products were confirmed by DNA sequencing.

### In vivo pulse-labeling analysis

*E. coli* MC1061 cells [12] transformed with the pING1 vector harboring the respective gene of interest were grown overnight at 37°C in M9 minimal medium supplemented with 19 amino acids (1 μg/ml) lacking Met, 100 μg/ml thiamine, 0.4% (w/v) fructose, 100 mg/ml ampicillin, 2 mM MgSO_4_, and 0.1mM CaCl_2_. Overnight cultures were diluted into fresh M9 medium to an OD_600_ of 0.1 and cell growth was continued until an OD_600_ of 0.3 was reached. Induction of protein expression was started by adding arabinose to a final concentration of 0.2% (w/v) and continued for 5 min at 37°C. After radiolabeling with [^35^S]-methionine for 2 min at 37°C the reaction was stopped by adding ice-cold trichloroacetic acid (TCA) to a final concentration of 10%. Proteins were precipitated on ice for 30 min and were spun down for 10 min at 20,000 g at 4°C. Samples were washed with ice-cold acetone, centrifugation was repeated and pellets were solubilized in Tris-SDS buffer (10 mM Tris-Cl pH 7.5, 2% (w/v) SDS) for 5 min at 1,400 rpm at 37°C. After another centrifugation step for 5 min at 20,000 g, the supernatant was incubated in TSET buffer (50 mM Tris-HCl pH 8.0, 150 mM NaCl, 0.1 mM EDTA-KOH, 2% (v/v) triton X-100) supplemented with Pansorbin^®^ for 15 min on ice. Proteins non-specifically bound to Pansorbin^®^ were removed by centrifugation at 20,000 g. The proteins of interest in the supernatant were further used for immunoprecipitation with Pansorbin^®^ and LepB antisera (in-house rabbit polyclonal IgG), which was carried out for at least 1 hour at 4°C whilst rolling. Immunoprecipitates were spun down and washed with 10 mM Tris-Cl pH 7.5, 150 mM NaCl, 2 mM EDTA, and 0.2% (v/v) triton X-100 and subsequently with 10 mM Tris-Cl pH 7.5. Pellets were solubilized in SDS sample buffer (67 mM Tris, 33% (w/v) SDS, 0.012% (w/v) bromophenol blue, 10 mM EDTA-KOH pH 8.0, 6.75% (v/v) glycerol, 100 mM DTT) for 10 min while shaking at 1,400 rpm. Solubilized proteins were incubated with 0.25 mg/ml RNase for 30 min at 37°C to hydrolyze tRNA and subsequently separated by SDS-PAGE. Gel fixation was done in 30% (v/v) methanol and 10% (v/v) acetic acid. After incubation in Gel-Dry^™^ Drying Solution for 30 min, gels were dried in a Bio-Rad gel dryer model 583. By exposing dried gels to phosphorimaging plates, radiolabeled proteins were detected using a Fuji FLA-3000 scanner. Band intensity profiles were obtained using the Fiji (ImageJ) software and quantified with the in-house software EasyQuant in order to determine the fraction full-length protein *f_FL_* = *I_FL_*/(*I_FL_* + *I_A_*), where *I_A_, I_FL_* are the intensities of the *A* and *FL* bands, respectively. *A_c_* and/or *FL_c_* size controls were included in the SDS-PAGE to confirm the identity of the *A* and *FL* bands. Each data point in the force profile represents the average of three independent biological replicates and includes the standard error of the mean (SEM).

## Results

### Force Profile Analysis

FPA takes advantage of the fact that a translational arrest peptide (AP) can induce a temporary pause in translation by binding in the upper parts of the ribosome exit tunnel when its last codon is positioned in the ribosomal A-site [13]. The AP-induced stalling can be overcome by pulling forces acting on the nascent chain [14,15], with different APs being sensitive to different force levels [16]. Such pulling forces can be generated by, *e.g*., cotranslational protein folding in the cytoplasm as well as in the periplasm, or cotranslational insertion of transmembrane segments into the membrane [9,17,18]. Therefore, APs can be conveniently used as force sensors to study cotranslational events *in vitro* and *in vivo*.

Here, we have used a series of constructs harboring a force-generating model TMH (H-segment) of composition *n*L/(19-*n*)A (*n* = 4, 6) placed *L* residues away from the C-terminal end of the 17-residue *E.coli* SecM (SecM(*Ec*)) AP followed by a 23-residue C-terminal tail [8], Figure 1a. The backbone of the design is from *E. coli* LepB, an inner membrane protein with two N-terminal TMHs and a periplasmic C-terminal domain [19]. The model 19-residue TMH of composition 4L/15A or 6L/13A (H-segment) followed by the SecM(*Ec*) AP is fused to the C terminus of the LepB part. A flanking region (FR), composed either of the positively charged loop between BtuC TMH1 and TMH2 (BtuC residues 47-59; Fig. 2a) or of the reentrant loop between BtuC TMH5 and TMH6 (BtuC residues 175-191; Fig 3a), was inserted immediately upstream of the H-segment (the amino acid sequences of all constructs are shown in Supplementary Fig. S1).

**Figure 1.**
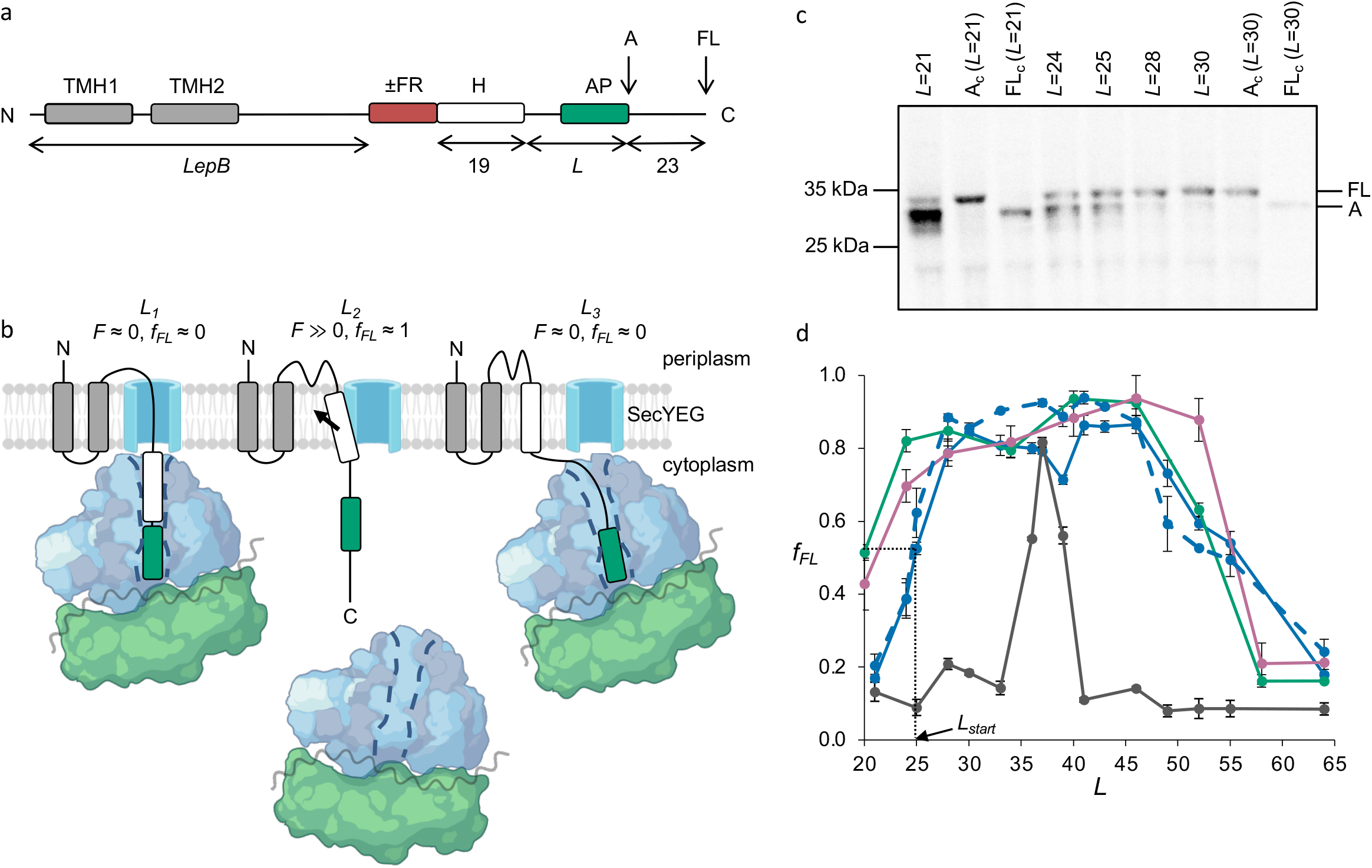
The force profile assay. (a) Basic construct. The LepB transmembrane helices TMH1 and TMH2, the flanking region (FR; positively charged loop or re-entrant loop), the H-segment, and the SecM(*Ec*) AP are indicated, together with the arrested (*A*) and full-length (*FL*) products. *L* denotes the number of residues between the C-terminal end of the H-region and the last residue in the AP. (b) At construct length *L_1_*, the H-region has not yet entered the SecYEG channel, no pulling force *F* is generated, and little full-length product is produced (*F* ≈ 0, *f_FL_* ≈ 0). At *L_2_*, the H-region is integrating into the membrane, generating a strong pulling force (*F* ≫0, *f_FL_* ≈ 1). At *L_3_*, the H-region is already integrated and little pulling force is generated (*F* ≈ 0, *f_FL_* ≈ 0). (c) SDS-PAGE gels showing *A* and *FL* products for [^35^S]-Met labelled and immunoprecipitated 4L/15A constructs with different *L*-values (c.f., panel *d*). Control constructs *A_C_* and *FL_c_* have, respectively, a stop codon and an inactivating Ala codon replacing the last Pro codon in the AP, leading to the production of *A*-sized and *FL*-sized proteins. Mw markers are indicated on the left. (d) FPs obtained with the SecM(*Ec*) AP for constructs lacking the FR segment and with H-segments composed of BtuC TMH2 (green), BtuC TMH6 (magenta), 4L/15A (blue), 6L/13A (dashed blue), and the 6L/13A construct but with the SecM(*Ec*-Sup1) AP (dark grey), c.f. [8,9]. The *L_start_* value for 4L/15A is indicated by the arrow. Error bars show SEM values (*n* = 3).

**Figure 2.**
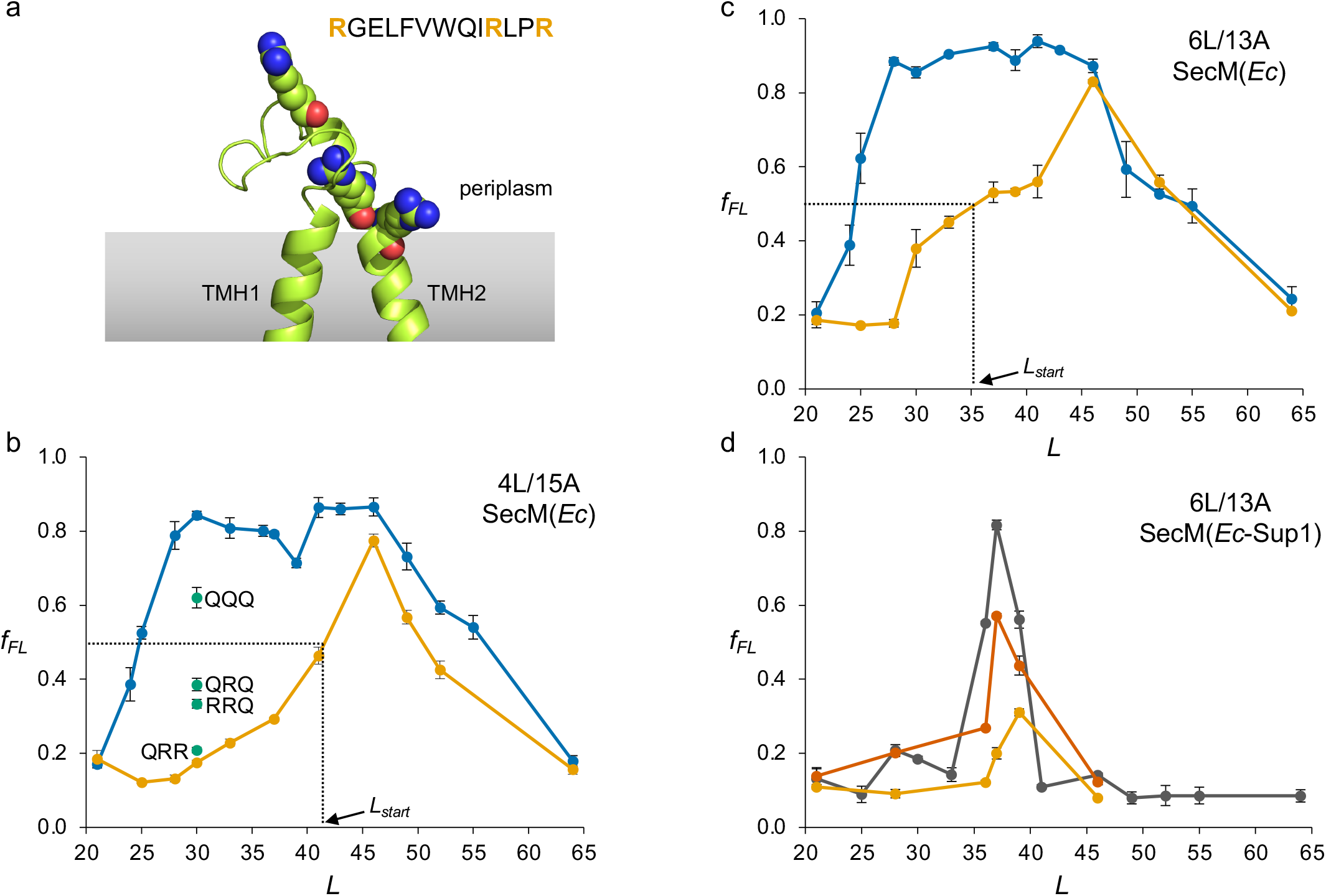
Analysis of the positively charged BtuC TMH1-TMH2 loop. (a) Structure of the BtuC TMH1-TMH2 region (PDB 2QI9) [27] with the three Arg residues in spacefill. The sequence of the loop is shown on top. (b) FPs for 4L/15A constructs lacking (blue) and including (orange) the positively charged loop (FR in Fig. 1a). Results for R→Q mutants are shown in green. (c) FPs for 6L/13A constructs lacking (blue) and including (orange) the positively charged loop (FR in Fig. 1a). (d) FPs obtained with the SecM(*Ec*-Sup1) AP for 6L/13A (dark grey), 6L/13A with the positively charged loop (orange), and 6L/13A with the re-entrant loop (red). Error bars show SEM values (*n* = 3).

**Figure 3.**
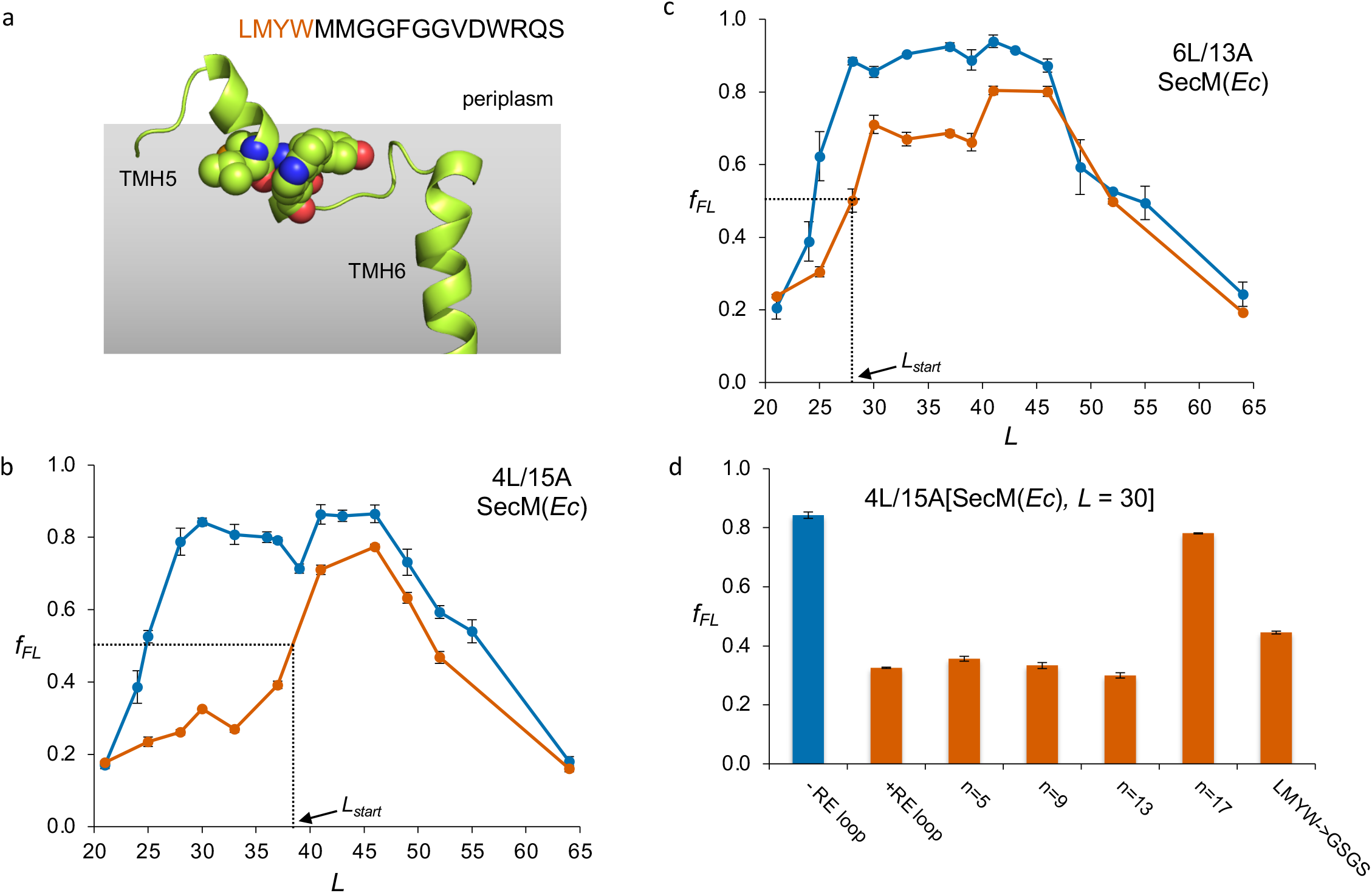
Analysis of the TMH5-TMH6 re-entrant loop. (a) Structure of the BtuC TMH5-TMH6 region (PDB 2QI9) [27] with the LMYW residues in spacefill. The sequence of the re-entrant loop is shown on top. (b) FPs for 4L/15A constructs lacking (blue) and including (red) the re-entrant loop (FR in Fig. 1a). (c) FPs for 6L/13A constructs lacking (blue) and including (red) the re-entrant loop (FR in Fig. 1a). (d) *f_FL_* values obtained with the SecM(*Ec*) AP for mutants of the re-entrant (RE) loop–4L/15A[L=30] construct where *n* = 5, 9, 13, and 17 residues in the re-entrant loop are replaced by GSGS repeats starting from the C-terminal end, or where the LMYW segment in the re-entrant loop is replaced by GSGS (see Supplementary Fig. S1 for sequences). The −RE and +RE values are from panel *b*. Error bars show SEM values (*n* = 3).

By varying *L*, the model TMH will be positioned in different locations in the ribosome exit tunnel or SecYEG translocon channel at the point when the ribosome reaches the last codon of the AP, and will therefore generate pulling forces of different magnitude. In a construct where membrane integration of the H-segment happens at the exact same time as the ribosome reaches the last codon of the AP, a strong pulling force *F* will be exerted on the nascent chain, which in turn prevents the translational pausing at the AP and results in mostly full-length protein (including the C-terminal tail) being produced during a short pulse with [^35^S]-Met, Figure 1b (middle). In contrast, in constructs where the H-segment has not yet reached the translocon, or has already been inserted into the membrane at the point when the ribosome reaches the end of the AP, little force is exerted on the AP, translational pausing will be efficient, and only the arrested form of the protein will be produced during a short pulse with [^35^S]-Met, Figure 1b (left and right). After immunoprecipitation with LepB antisera, full-length (*FL*) and arrested (*A*) species are separated on an SDS-PAGE gel and the fraction full-length protein is calculated, *f_FL_* = *I_FL_*/(*I_FL_*+*I_A_*), where *I_FL_* and *I_A_* are the intensities of the bands representing *FL* and *A* species, respectively (Figure 1c). Because it is sensitive to pulling force,*f_FL_* can be used as a proxy for *F* [15,20–22]. A force profile (FP), in which *f_FL_* is plotted against *L* then can be used to follow, step-by-step, the membrane integration of model or natural TMHs; FPs for the 4L/15A and 6L/13A H-segments as well as for BtuC TMH2 and TMH6 are shown in Fig. 1d (see Supplementary Fig. S1 for amino acid sequences of all constructs). For a given peak in the FP, we define *Lstart* as the *L* value for which *f_FL_* reaches its half-maximal value (*L_start_* ≈ 25 residues for the 4L/15A and 6L/13A H-segments, *L_start_* ≈ 22 residues for BtuC TMH2 and TMH6, Fig. 1d)

According to the ΔG predictor [4], BtuC TMH2 and TMH6 are predicted to form TMHs with ΔG values of −0.13 and −0.79 kcal/mol, respectively, and both generate strong pulling force from *L*=24 to *L*=52 [9], Fig 1d. The model H-segments 4L/15A and 6L/13A [8] were chosen based on similar ΔG values of −0.49 and −1.54 kcal/mol, respectively. As expected, the model and natural TMHs have very similar FPs, with slightly earlier *L_start_* values for the latter. An FP was also obtained for the 6L/13A H-segment using the stronger AP variant SecM(*Ec*-Sup1) [23], showing that the maximal force is exerted at *L_max_* = 37 residues, Fig 1d.

### Upstream Arg residues delay membrane insertion of a model TMH

In our recent FPA study of the cotranslational membrane insertion of BtuC, we observed that TMH2 initiated membrane insertion when its N-terminal end is ~55 rather than the customary ~45 residues away from the PTC [9], and provisionally attributed this unexpected finding to the presence of the highly positively charged periplasmic loop between TMH1 and TMH2, Fig. 2a. According to the ‘positive-inside rule’ [24], short periplasmic loops between TMHs typically contain only 0-2 positively charged residues, and the TMH1-TMH2 loop that contains 3 Arg residues is thus an unusual feature in bacterial inner membrane proteins.

In order to simplify the experimental system and focus on the possible effects on membrane insertion of the positively charged loop, we used the construct design shown in Fig. 1a. To obtain the FP for the membrane insertion of the H-segment, the length *L* of the linker between the H-segment and the AP was increased stepwise from *L* = 21 to *L* = 64 residues, and *f_FL_* was measured for each construct. As shown in Fig. 2b-c, in the absence of the positively charged loop, the 4L/15A and 6L/13A TMHs exert half-maximal pulling force at *L_start_* ≈ 25 residues, *i.e*., when their N-terminal ends are 25+19 = 44 residues from the PTC, as expected. When preceded by the positively charged loop, however, half-maximal pulling force is reached only at *L_start_* ≈ 41 residues for 4L/15A and *L* ≈ 35 residues for 6L/13A. Indeed, *f_FL_* remains at its background value until *L* ≈ 30 for both H-segments in the presence of the positively charged loop. Using the much stronger SecM(*Ec*-Sup1) AP [23], we see that not only is membrane insertion of the 6L/13A H-segment delayed by the positively charged loop, but the maximal pulling force exerted during the insertion process is also reduced, Fig. 2d (orange curve).

To better appreciate the role of the three different Arg residues, we made four R→Q mutants in the 4L/15A[L=30] construct (green dots in Fig.2b). The R^47^→Q mutation (QRR) had almost no effect on *f_FL_*, while the R^59^→Q mutation, either alone (RRQ) or together with R^47^→Q (QRQ), gave rise to a substantial increase in *f_FL_*. A very strong effect on *f_FL_* was seen when all three Arg residues, including R^56^, were mutated (QQQ). Thus, the main culprits in the positively charged loop are R^56^ and R^59^, *i.e*., the two Arg residue located closest to the H-segment. We conclude that positively charged residues located close to the N terminus of a Nout-oriented TMH can substantially delay its insertion into the membrane.

### A short upstream hydrophobic segment delays membrane insertion of a model TMH

In our recent study of the membrane insertion of BtuC [9], we also found that the insertion of TMH6 gave rise to two peaks in the FP, one that appeared when the N-terminal end of TMH6 was only 37 residues away from the PTC and one when it was 63 residues away, and we provisionally attributed these unexpected observations to the presence of an upstream reentrant loop located between TMH5 and TMH6 (Fig. 3a) and an amphipathic surface helix directly downstream of TMH6. As seen in Fig. 1d, when TMH6 (BtuC residues 175-191) is analyzed in the context of the LepB construct (Fig. 1a), half-maximal pulling force is seen at *L_start_* ≈ 22 residues, *i.e*., when the N-terminal end of TMH6 is 41 residues from the PTC, similar to the results for the 4L15A and 6L/13A H-segments.

To directly probe the role of the re-entrant loop (BtuC residues 175-191) we inserted it upstream of the 4L/15A and 6L/13A H-segments (‘FR’ in Fig 1a) and recorded the respective FPs, Fig. 3b-c. For the 4L/15A series of constructs, *L_start_* ≈ 38 residues, *i.e*., the re-entrant loop delays the insertion of the H-segment by ~13 residues. A smaller delay of ~3 residues is seen with the more hydrophobic 6L/13A H-segment, Fig 3c. The effect of the re-entrant loop on the insertion of TMH6 seen in the intact BtuC protein can thus be reproduced with the model H-segments. As for the positively charged loop, an FP obtained with the stronger SecM(*Ec*-Sup1) AP shows both slightly delayed membrane insertion of the 6L/13A H-segment and a reduction in the maximal pulling force exerted during the insertion process in the presence of the re-entrant loop, Fig. 2d (red curve).

To pin-point the critical residues in the re-entrant loop, we replaced an increasing number of loop residues in the 4L/15A[*L*=30] construct, first five and then four at a time, by GS-repeats, starting from the C-terminal end of the loop, Fig. 3d. Thirteen loop residues could be replaced by GS repeats with no effect on *f_FL_*; only when the entire re-entrant loop segment was replaced by GS repeats did *f_FL_* return to the high value seen for the 4L/15A H-segment itself. The four most N-terminal residues in the re-entrant loop (LMYW) thus seem to be the main determinants of the delay in membrane insertion of the H-segment and, indeed, when these four residues were mutated to GSGS while keeping the other residues in the re-entrant loop there was a clear increase in *f_FL_* compared to re-entrant loop construct (+RE loop), albeit not as strong as for the *n* = 17 mutant. The re-entrant loop, and especially the short hydrophobic helix at its N-terminal end, thus causes a marked reduction of the *f_FL_* values in the early part of the FP not only for TMH6 in BtuC but also for the model 4L/15A and 6L/13A H-segments in the LepB constructs. We conclude that a short hydrophobic stretch located ~15 residues upstream of the N terminus of a N_out_-oriented TMH can substantially modify the behavior of the TMH during the membrane insertion process.

## Discussion

The ‘sliding’ models for SecYEG-mediated cotranslational insertion of TMHs into the *E. coli* inner membrane posits that the hydrophobic segment makes contact with membrane lipids through a more or less open lateral gate as soon as it enters the SecYEG channel [25]. The current best estimate is that this happens when the N-terminal end of the hydrophobic segment is ~45 residues away from the PTC [7,9]. However, our recent study of the cotranslational membrane insertion of three multi-spanning *E. coli* inner membrane proteins [9] uncovered a couple of instances where membrane insertion of a particular TMH appeared to be delayed to markedly longer nascent chain lengths. In order to better understand these exceptional cases, we have tried to reproduce them in a simplified system where short loop segments presumed to be responsible for the delay are inserted upstream of model TMHs (H-segments) composed only of Leu and Ala residues.

Our results show that positively charged Arg residues located close to the N terminus of an N_out_-oriented H-segment appear to completely prevent the latter from making hydrophobic interactions during an early stage of passage into the SecYEG channel, as seen in the FPs in Fig. 2b-c (*L* = 20-30 residues). A rough estimate suggests that this early stage corresponds to a situation where the Arg residues are near the hydrophobic gasket [10] that seals off the cytoplasmic from the periplasmic parts of the channel, Fig. 4a. Once a sufficiently long stretch of the H-segment has entered the channel (*L* > 30 residues) there is an increase in *f_FL_*, indicating that the H-segment now engages in hydrophobic interactions and starts to insert into the membrane. Even in these later stages, however, the positively charged residues cause a persistent reduction in the *f_FL_* values. We propose that the electrical membrane potential pushes back on the positively charged residues and prevents them from translocating across the hydrophobic gasket (in turn preventing the H-segment from entering the SecYEG channel) until a sufficiently long segment of the H-segment has emerged from the ribosome exit tunnel such that the free energy gained upon its membrane insertion compensates for the energetic cost of moving the Arg residues across the gasket. The reduction in pulling force caused by positively charged residues is a mirror image of the increase in pulling force seen when negatively charged residues translocate across the SecYEG channel [21].

**Figure 4.**
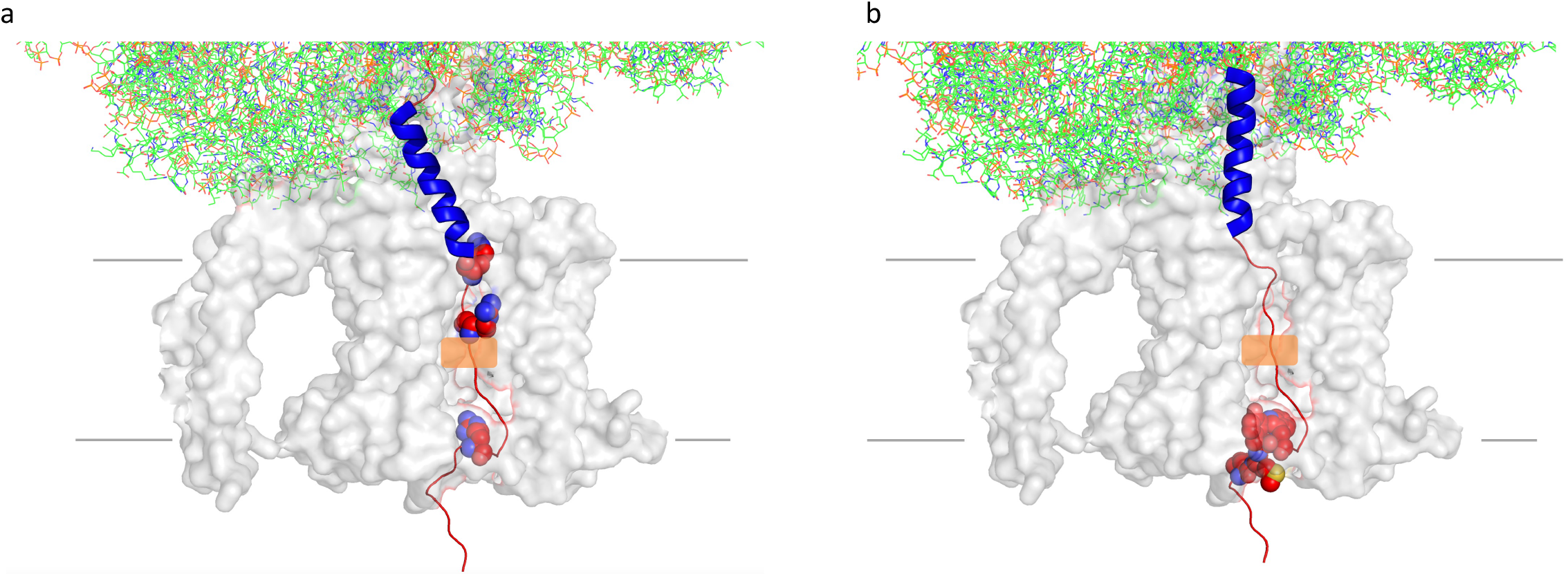
Models of ribosome-nascent chain-SecYEG complexes. (a) Model for the positively-charged loop *L* = 28 construct with the H-segment in blue and the three Arg residues in the positively charged loop in spacefill. The location of the hydrophobic gasket is indicated in orange. The model is based on PDB 4V6M [28], assuming that the H-segment is helical [29–32] and that the nascent chain model in the ribosome exit tunnel in the PDB file is stretched by ~2 residues compared to the SecM AP [33]. (b) Same as in panel *a*, but for the re-entrant loop *L* = 25 construct. The LMYW residues are in spacefill.

In contrast to the effect on insertion caused by the positively charged residues, the re-entrant loop does not reduce *f_FL_* to background values in the early part of the FP, but only attenuates the *f_FL_* values in this region, Fig. 3b-c. A rough model of the nascent chain at *L* = 25 residues, Fig. 4b, suggests that at this point the critical hydrophobic segment in the re-entrant loop (LMYW) has just reached the periplasmic exit from the SecYEG channel and is able to partition into the periplasmic leaflet of the membrane, while the TMH is just entering the translocon channel. We hypothesize that, when in this location, the re-entrant loop interacts with the lateral gate in such a way that it shifts the equilibrium towards the closed state [26] and thereby reduces the interaction between the TMH and the membrane.

In summary, we find that both charged and hydrophobic upstream segments can impact the timing of cotranslational insertion of model TMHs into the *E. coli* inner membrane, presumably reflecting both the transmembrane electric field and the dynamics of the lateral gate in the SecYEG channel.

## Acknowledgments

We thank Dr. Rickard Hedman (Stockholm University) for programming and maintenance of the EasyQuant software, and for providing the 4L/15A SecM(*Ec*), 6L/13A SecM(*Ec*) and 6L/13A SecM(*Ec*-Sup1) constructs used in this study. This work was supported by grants from the Knut and Alice Wallenberg Foundation (2017.0323), the Novo Nordisk Fund (NNF18OC0032828), and the Swedish Research Council (621-2014-3713) to GvH, and by a Marie Curie Initial Training Network Grant (Horizon 2020, ProteinFactory 642863) to FN.

---

**Supplementary Figure S1.**
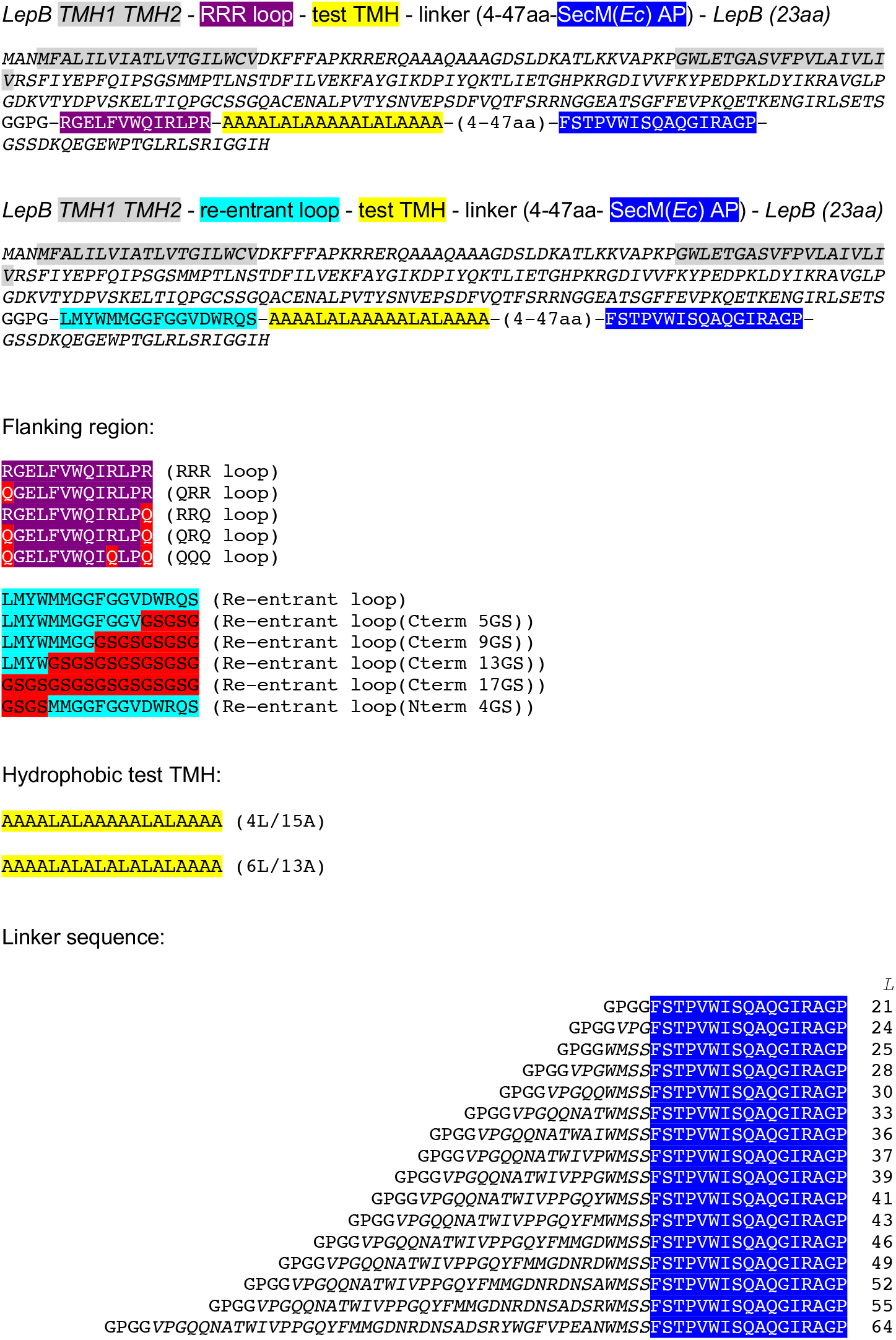
Design of protein sequences

